# Multiscale ATUM-FIB microscopy enables targeted ultrastructural analysis at isotropic resolution

**DOI:** 10.1101/2020.03.30.015727

**Authors:** Georg Kislinger, Helmut Gnägi, Martin Kerschensteiner, Mikael Simons, Thomas Misgeld, Martina Schifferer

## Abstract

Volume electron microscopy enables the ultrastructural analysis of biological tissues and is essential for dense reconstructions e.g. of neuronal circuits. So far, three-dimensional analysis is based on either serial sectioning followed by sequential imaging (ATUM, ssTEM/SEM) or serial block-face imaging (SB-SEM, FIB-SEM), where imaging is intercalated with sectioning. Currently, the techniques involving ultramicrotomy allow scanning large fields of view, but afford only limited z-resolution determined by section thickness, while ion beam-milling approaches yield isotropic voxels, but are restricted in volume size. Now we present a hybrid method, named ATUM-FIB, which combines the advantages of both approaches: ATUM-FIB is based on serial sectioning of tissue into semithick (2-10 µm) resin sections that are collected onto transparent tape. 3D information obtained by serial light and electron microscopy allows identifying regions of interest that are then directly accessible for targeted FIB-SEM. The set of serial semithin sections thus represent a tissue ‘library’, which provides information about microscopic tissue context that can then be probed ‘on demand’ by local high resolution analysis. We demonstrate the potential of this technique to reveal the ultrastructure of rare but pathologically important events by identifying microglia contact sites with amyloid plaques in a mouse model for familial Alzheimer’s disease.

## Introduction

Since the completion of the first connectomics data set (White, Southgate, Thomson, & Brenner, 1986) volume EM techniques have been substantially refined and advanced. While the interest in deciphering neuronal networks was the major driving force behind these technological developments, three-dimensional (3D) ultrastructural analysis has attracted considerable recent attention in a wide range of biological fields (Titze & Genoud, 2016). Applications range from classical cell biological questions to developmental, neuro- and cancer biology to microbiology and botany (Karreman, 2014). Currently, 3D ultrastructure can be solved by destructive techniques including serial block-face electron microscopy (SB-SEM) (Briggman, Helmstaedter, & Denk, 2011; Denk & Horstmann, 2004; Helmstaedter et al., 2013; Mikula & Denk, 2015) or focused ion beam-scanning electron microscopy (FIB-SEM) (Heymann et al., 2006; Knott, Marchman, Wall, & Lich, 2008; Sonomura et al., 2013). While these methods benefit from high alignment accuracy, they lack the option of reacquisition and hierarchical imaging (Kornfeld & Denk, 2018). Alternatively, serial sectioning for transmission microscope camera array (TEMCA (Bock et al., 2011; Lee et al., 2016; Zheng et al., 2018) or automated tape-collecting ultramicrotomy (ATUM) (K. J. Hayworth et al., 2014; Hildebrand et al., 2017; Kasthuri et al., 2015; Mikula & Denk, 2015; Morgan, Berger, Wetzel, & Lichtman, 2016; Schalek et al., 2011; Terasaki et al., 2013; Tomassy et al., 2014) allow repetitive acquisition of the same or other regions of interest. However, as ultramicrotomy-based approaches are limited by their poor z resolution of maximally 30 nm, ion milling techniques are required, if isotropic high resolution voxels are needed. Still, the imaging volume in FIB-SEM is limited to a few tens of microns due to the accumulation of milling artifacts caused by high-energy gallium ions (Xu et al., 2017). Despite recent advances in the application of alternative milling strategies (Kornfeld & Denk, 2018), targeted imaging is still required to restrict the acquisition volume. This is mainly achieved by correlated workflows involving targeted trimming guided by endogenous (Luckner et al., 2018) and artificial landmarks (Bishop et al., 2011; M. A. Karreman, Hyenne, Schwab, & Goetz, 2016; Matthia A. Karreman et al., 2014; Villani et al., 2019). X-ray micro computed tomography (microCT) (Bushong et al., 2015; Matthia A. Karreman et al., 2014; Sengle, Tufa, Sakai, Zulliger, & Keene, 2012; Villani et al., 2019) has emerged as a tool for facilitated ROI relocation within the processed EM sample. This not only bridges multiple scales, from millimeter to micrometer dimensions, but also puts the site of interest into a wider morphological context (Maire & Withers, 2014). However so far, microCT imaging options are not commonly accessible and the technique only provides a virtual map for subsequent guided destructive sample preparation. An alternative prescreening of embedded tissue at a larger scale is implemented by rendering it accessible to light and electron microscopy (EM) modalities. Ultrathick sectioning at 20 µm by the hot knife method provides samples that are accessible to large-scale FIB-SEM (Kenneth J. Hayworth et al., 2015) enabling seamless reconstruction of large tissue blocks. While this method was designed for a complete reconstruction of big volumes, it would be desirable to reduce time and data load for biological questions requiring targeted ultrastructural analysis.

Here, we developed a multiscale method for targeted FIB-SEM on semithick (2-10 µm) sections named ATUM-FIB that combines the advantages of both ultramicrotomy-based and serial block face imaging approaches. This approach is based on the ultramicrotomy on partially cured (Droz, Rambourg, & Koenig, 1975) resin-embedded samples facilitated by a custom-built diamond knife with temperature control. Serial thick sections are then collected onto carbon nanotube (CNT) tape (Kubota et al., 2018), compatible with both serial bright-field and scanning electron microscopy (SEM). While providing direct physical access to isotropic high-resolution imaging of multiple ROIs by FIB-SEM, this method at the same time provides 3D tissue context information and allows archiving samples for future extended analysis.

## Results

### Semithick sections provide suitable information content for sparse ultramicrotomy-guided targeting

Here, we developed a method that allows the search for an ultrastructural feature within a volume and the subsequent acquisition of a defined isotropic volume by FIB-SEM (Fig. 1). Our goal was to expose surfaces at defined distances by serial sectioning. While traditional ultrathin sectioning combined with sparse imaging saves imaging time, it is not compatible with the investigation of isotropic volumes. In order to overcome this limitation and optimize ultramicrotomy time as well as screening efficiency, the information content for FIB-SEM increases with section thickness, we therefore explored whether a “semithick” sections ranging from 0.5 to 10 µm would meet these requirements (Fig. 2A). Moreover, we aimed to apply water instead of an oil bath (as used in the ultrathick partitioning approach (Kenneth J. Hayworth et al., 2015)) in order to avoid additional re-embedding and sectioning.

**Figure 1.**
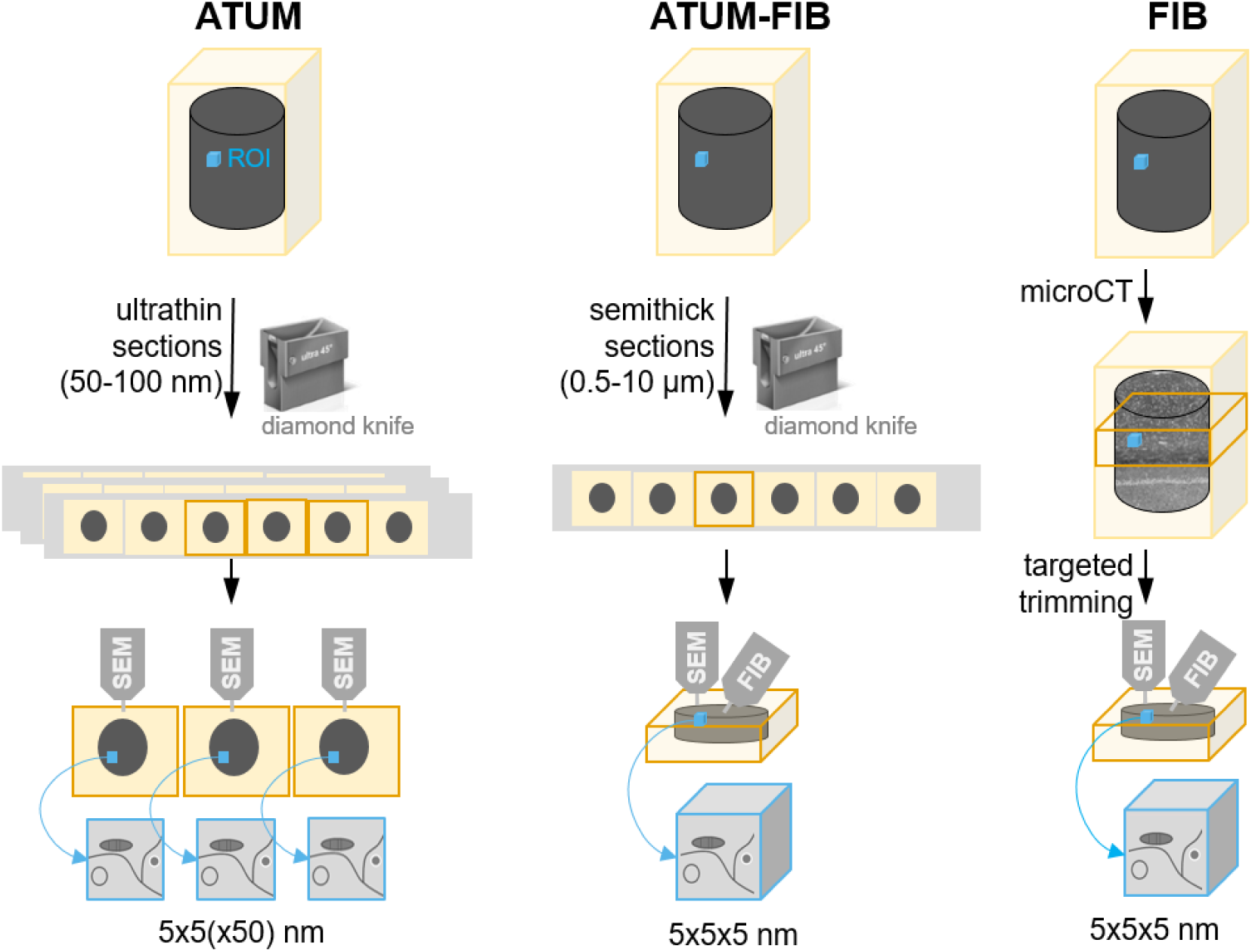
Principle of ATUM-FIB as s a hybrid volume EM approach. Schematic of existing techniques for targeted volume SEM and comparison to the new ATUM-FIB approach (middle). Both, ATUM and ATUM-FIB are microtomy-based, but ATUM-FIB generates semithick sections that can be selected by serial section light microscopy and SEM and subjected to further FIB-SEM investigation.

**Figure 2.**
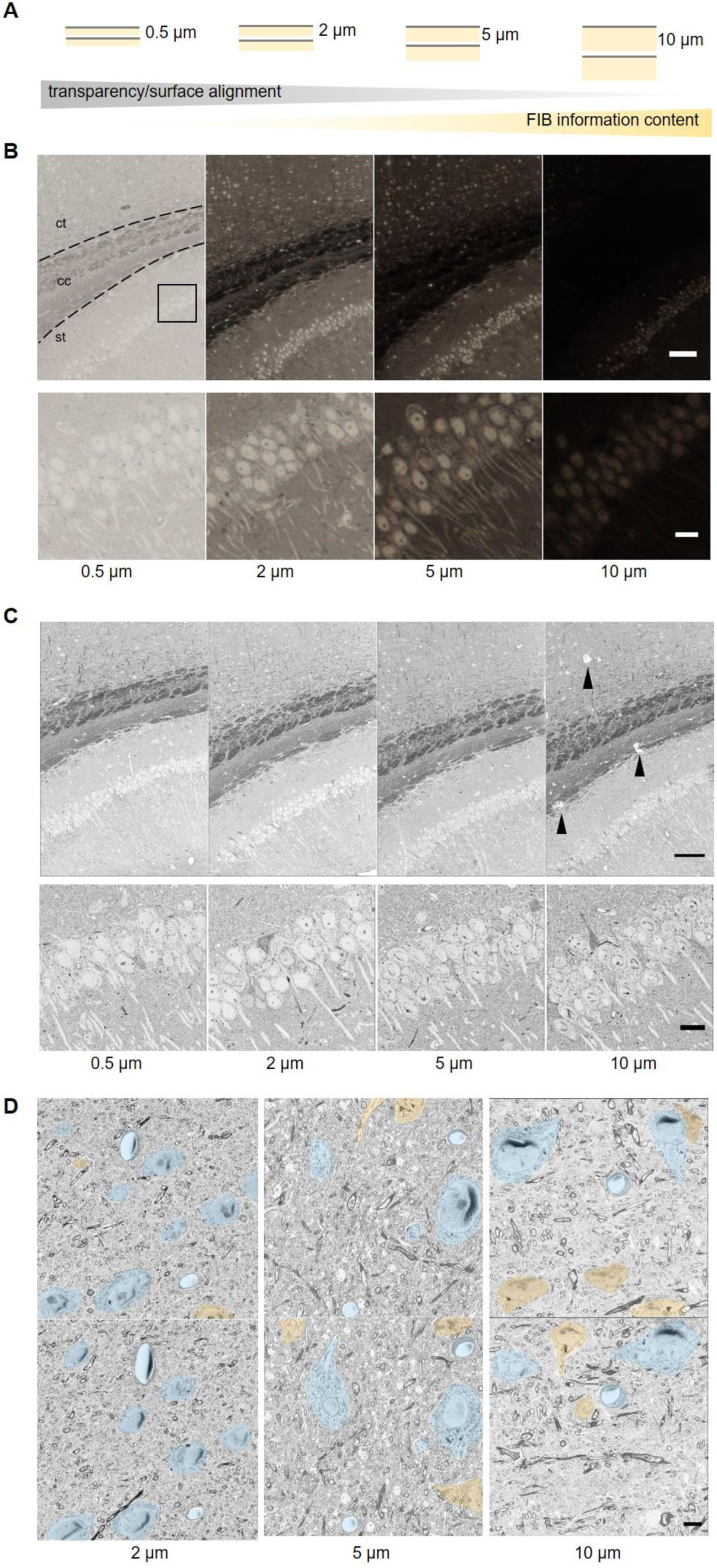
Targeting regions of interest by semithick sectioning. (A) Cartoon of the information content of BSD surface imaging and the potential FIB-SEM analysis of sections of different thickness. (B) Transmitted light and (C) BSD images of 0.5, 2, 5 and 10 µm sections of mouse brain tissue. Corpus callosum (cc), cortex (ct), striatum (st). Scale bars 100 µm (top), 10 µm (bottom). (D) Consecutive (upper, lower row) semithick (2, 5, 10 µm) sections allow tracing cell bodies and blood vessels across sections. Somata and blood vessels that can be followed from one to the next section are colored blue, others are shown in orange. Scale bar 5 µm.

Semithick sections (0.5-10 µm) were generated using a diamond knife. Carbon nanotube (CNT) tape was chosen as a conductive and transparent section support for sequential light and electron microscopic assessment (Kubota et al., 2018). The wet tape with sections was placed onto a glass slide on a heating plate (80°C) for better adherence and imaged on a slide scanner (Fig. 2B). Transmitted light investigation was possible for sections below 5 µm as reduced transparency masked morphological details. The sections were transferred onto a silicon wafer and the surface scanned by backscattered detection SEM which enables the exploration of the entire semithick section range (Fig. 2C). We generated two consecutive sections with 2, 5 and 10 µm thickness and looked at traceability of neuronal cell bodies and blood vessels. While most of the structures could be recognized on the next 2 µm section, alignment between two sections was difficult at thicknesses above 5 µm (Fig. 2D). In summary, combined light microscopy and SEM with maximal FIB-SEM information content is optimal at 2 µm while 5 µm sections can optimally be assessed by SEM surface imaging alone while conserving alignment of biological structures like cell bodies.

### Sequential resin curing and heating enables semithick sectioning and FIB milling on the same sample

The generation of resin sections thicker than 2 µm necessitated the optimization of resin characteristics and ultramicrotomy parameters. We therefore explored different contrasting methods and resins for semithick sectioning and subsequent SEM imaging. First, for contrasting, we settled on a standard rOTO (reduced osmium thiocarbohydrazide osmium) protocol without lead aspartate impregnation (Hua, Laserstein, & Helmstaedter, 2015), as post-contrasting strategies, e.g. incubation in uranylacetate in ethanol at 60°C would risk uneven stain distribution within and between thick sections (Kenneth J. Hayworth et al., 2015),(Locke & Krishnan, 1971). Second, we optimized the resin choice. Resin requirements for thick sectioning and ion milling are conflicting: For semithick sectioning, softer resin and less heavy metal staining would be favoured, while FIB-SEM requires hard resin and good contrasting to avoid charging artefacts. Various methacrylate (Norris, Baena, & Terasaki, 2017) but mainly epoxy resins including epon and durcupan (Kenneth J. Hayworth et al., 2015) have been used for standard ultrathin serial sectioning. We investigated rOTO-processed mouse cortex samples embedded in different resins by semithick sectioning and subsequent backscattered detection SEM (Fig. 3A). Regarding the sectioning characteristics, epon formulations as well as durcupan - which would be preferable for FIB-SEM - showed very uneven surface topology at thicknesses above 1 µm (Fig. 3A). Tissue embedded in epoxy resin LX112-embedded tissue could be sectioned up to 2 µm without major surface defects (Fig. 3A). We reasoned that a two-step curing (Droz et al., 1975) would yield in resin characteristics that are optimal for both, sectioning and FIB milling. Tissue embedded in LX112 was precured at 60°C for 10 and 48h. Semithick sections of up to 10 µm could be generated from the 10h cured blocks (Fig. 3A). Resin blocks were too soft and sticky for sectioning after shorter curing periods (data not shown). After ultramicrotomy, sections were post-cured for 2d at 60°C ensuring beam resistance required for both serial SEM and FIB-SEM imaging. Still, semithick sectioning proved difficult even in this softer pre-cured resin, there for we explored a ‘hot knife’ approach (Kenneth J. Hayworth et al., 2015).

**Figure 3.**
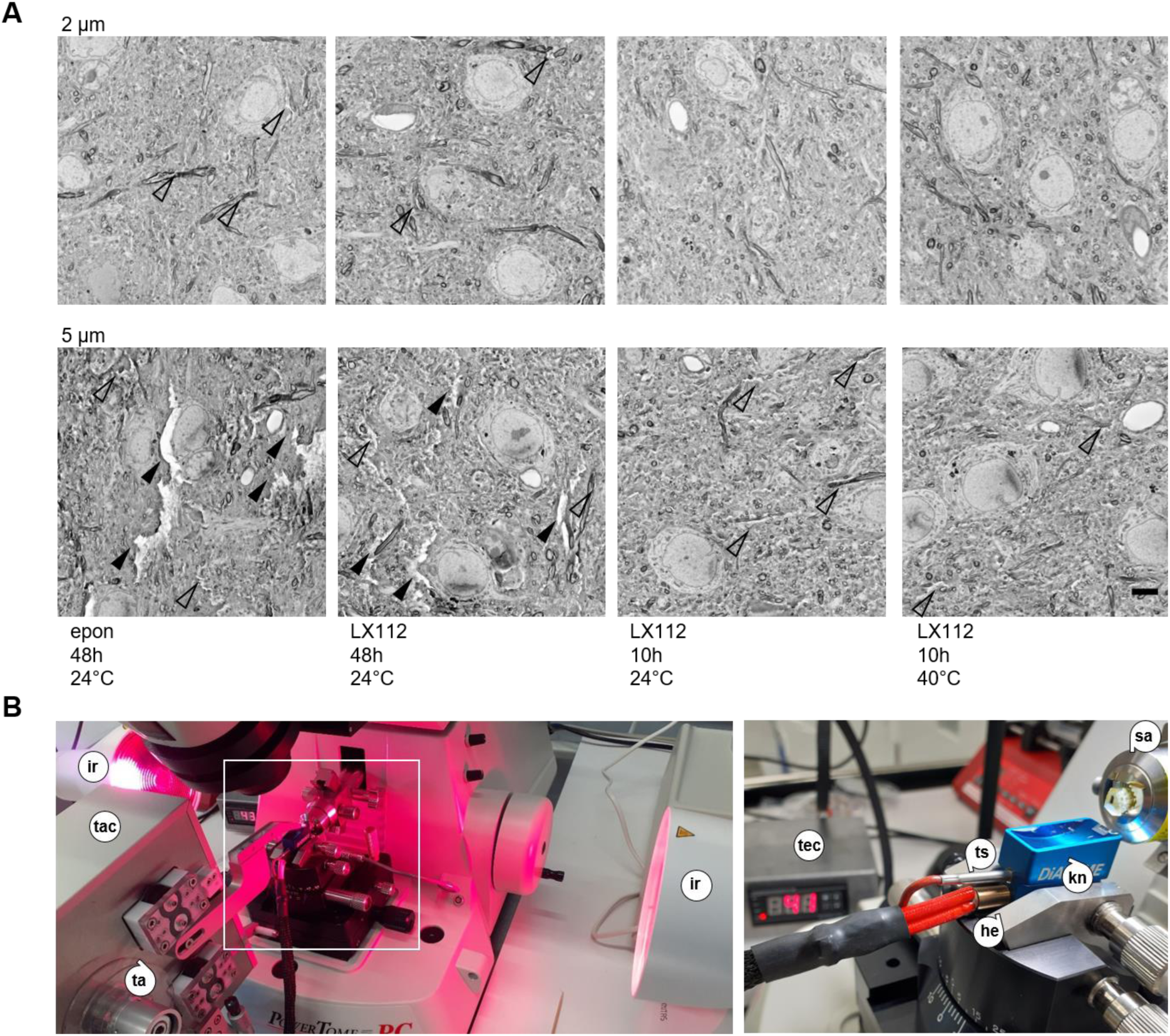
Generation of semithick sections by sequential curing and elevated-temperature microtomy. (A) Comparison of resin performance for semithick sections of mouse cortex tissue. BSD images were acquired of 2 (top) and 5 µm (bottom) semithick sections of epon and LX112 blocks cured for 48 or 10h and sectioned at room temperature or at 40°C. Left to right: epon cured for 48h, sectioned at 24°C; LX112 cured for 48h, sectioned at 24°C; LX112 cured for 10h, sectioned at 24°C; LX112 cured for 48h, sectioned at 40°C. Scale bar 5 µm. Arrowheads indicate major (full) and minor (open) tissue cracks. (B) Photographs of infrared lights installed at the sides of the microtome sample arm to warm the sample during semithick sectioning (left). The heated diamond knife is depicted with a lower heater and the upper sensor sticks and the temperature control (right). Heating element (he), infrared light (ir), sample arm (sa), tape (ta), tape collector (tac), temperature control (tec), temperature sensor (ts).

In order to facilitate semithick sectioning, we developed a heated diamond knife (Fig. 3B). We drilled holes into the 35° and 45° ultra boats for a temperature sensor (3 mm diameter) and a heater (6 mm diameter). For further temperature stability of the sample itself we installed infrared lights at both sides and adjusted the distance to the ultramicrotome sample arm to yield a sample temperature of 40 °C (Fig. 3B). At temperatures above 40°C, water evaporated onto the sample block which partially melted (data not shown). We compared sectioning of rOTO processed mouse cortex precured for 10h in LX112 at 0.5-10 µm thickness at room temperature, 30°C and 40°C. Higher temperatures increased the smoothness of the section surface (Fig. 3A). At higher temperatures, the water pumping system supplying the knife boat had to be adapted to increased flow rates to compensate increased evaporation. Therefore, a good compromise for serial sectioning were temperatures in the range of 35-40°C. Comparison of 35° ultra- with 45° ultra knives didn’t show major differences in semithick sectioning (data not shown), but higher long-term robustness is expected for the 45° knife (Matzelle, Gnaegi, Ricker, & Reichelt, 2003). Consequently, a combination of two-step curing of LX112 and heated ultramicrotomy allows for semithick sectioning at 0.5-10 µm.

### Light and electron microscopy of serial semithick sectioning reveals ultrastructural details for selection for FIB-SEM imaging

For volume analysis of cortical mouse tissue we collected serial semithick sections on CNT tape using a tape collector (Powertome, RMC) (Fig. 4A). Sections were collected at 0.2-0.3 mm/sec with increased tape speed within and reduced tape speed outside the cutting window to assure the efficient and compact uptake. Slow speed was needed both for limiting compression and in order to keep the section longer on the heated water bath to smoothen. For test purposes, we cut 50 sections of mouse cortex tissue at 5 µm. After sectioning and collection, the CNT tape was cut into 5 cm strips and reversibly adhered onto glass slides by application of a drop of water and incubation at 60° on a heating plate. For 5 µm thick sections we acquired section overview images by transmitted light microscopy (Fig. 4A). Samples were mounted onto a silicon wafer as previously described (Djannatian et al., 2019; Kasthuri et al., 2015) and postcured. BSD images were captured at 4kV (Fig. 4B). Serial section images were taken at 0.2 × 0.2 × 5 µm and regions of interest at 0.01 × 0.01 µm. The beam dwell time was reduced to values below 2 µs/pixel in order to avoid charging artefacts originating from increased sample thickness. Some chargin on blood vessels or nuclei could not be avoided. Maximum projections of mouse cortex revealed morphological features including blood vessels (Fig. 4B).

**Figure 4.**
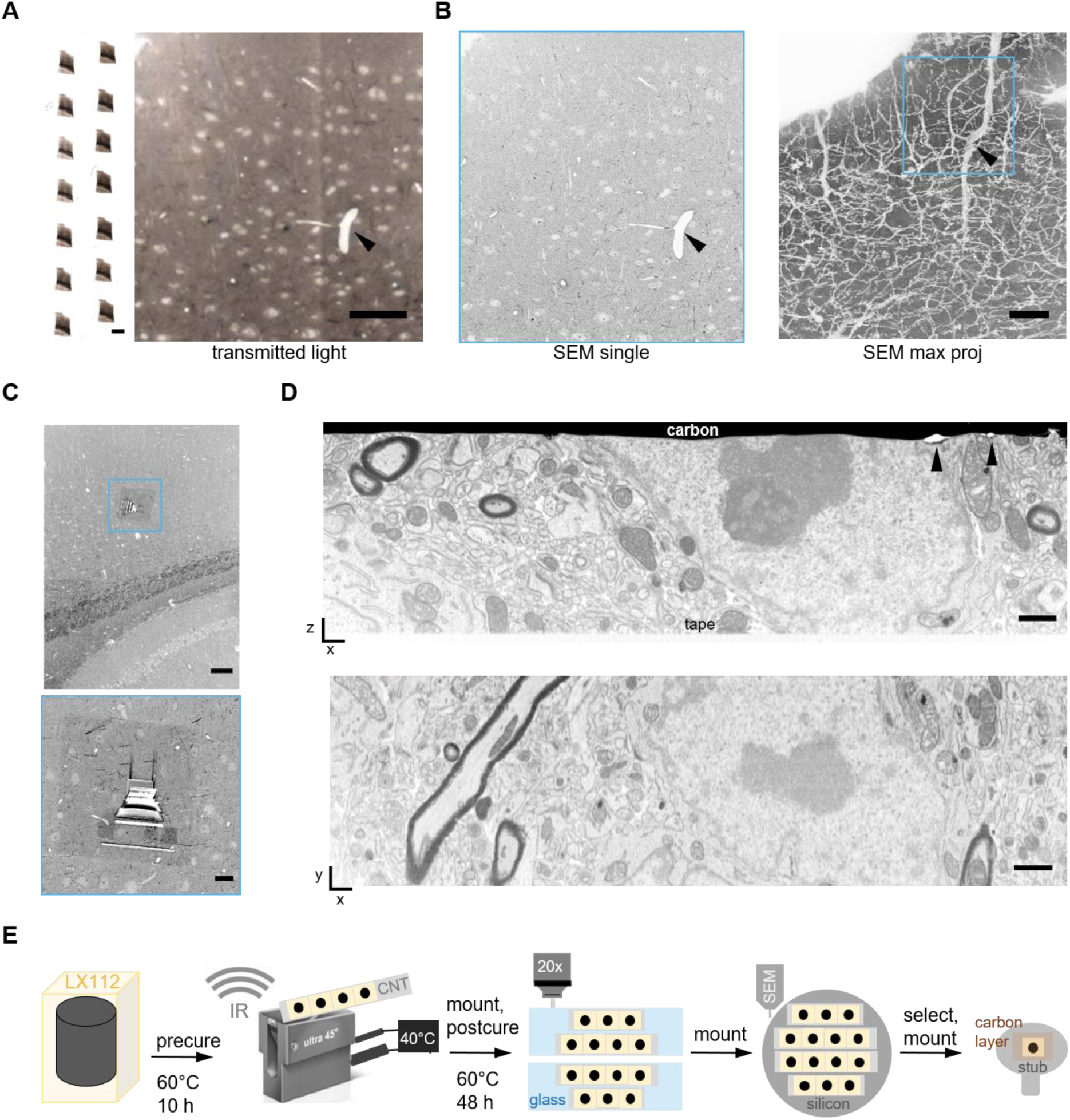
Serial semithick sectioning for targeted FIB-SEM. Serial 5 µm sections of mouse cortex were collected on CNT tape. (A) Overview of several consecutive sections on tape (left) and a single transmitted light micrograph (20x objective, right). Scale bars 2 mm and 100 µm, respectively. (B) BSD image (10×10×5000 nm) of one section (left) and maximum projection of all 50 sections showing blood vessel morphology (right). (C) A random cortical FIB target site was chosen (blue box) (top). The selected region was prepared for FIB-SEM by carbon deposition and trench milling (bottom). Scale bars 100 and 10 µm, respectively. (D) Cross section after FIB-SEM preparation showing the section – CNT tape adhesion site and the surface covered by a carbon layer (top). Defects in the surface layer topology are highlighted (arrowheads). After the FIB-SEM run a xy section was reconstructed (bottom) from 2000 SE images (resolution 5×5×5 nm^3^, bottom). Scale 1 µm. (E) Schematic of the ATUM-FIB strategy: sequential resin (LX112) curing and heated microtomy are required for semithick sectioning. Serial sections are attached onto glass slides for light microscopy and remounted onto silicon wafers for serial SEM imaging and target selection. A section of interested is remounted, adhered onto a stub and carbon-coated for FIB-SEM examination.

Mouse cortical sections with 5 µm thickness were detached from the wafer and mounted onto a FIB-SEM stub by conducting carbon cement. A fine carbon layer was sputtered onto the sample providing surface accessibility for the electron beam while increasing conductivity. FIB-SEM was performed by milling a 8 µm deep trench (Fig. 4C). We acquired a FIB-SEM of a 30 × 30 × 5 µm cortical region data at 5 × 5 × 5 nm (Fig. 4D). Resin postcuring resulted in good milling characteristics without curtaining effect or other problems arising from samples embedded in soft epon. In summary, semithick sections can be imaged by light or electron microscopy to reveal (ultrastructural) details required for target site selection (Fig. 4E). Selected regions on a particular section can be subjected to isotropic voxel acquisition by FIB-SEM.

### Targeted FIB-SEM for isotropic ultrastructural analysis of amyloid plaques in FAD

As a proof of concept we combined serial semithick sectioning with targeted FIB-SEM to analyze microglia contacting dystrophic neurites in the cortex of a familial AD mouse model. Familial Alzheimer’s Disease (AD), the most common form of dementia in the elderly, causes gradual loss of memory, judgment and the ability to function socially. It is characterized by the presence of extracellular plaques composed of amyloid-β (Aβ) peptides and intracellular tau aggregates (Giacobini & Gold, 2013; Kwak et al., 2020). The plaques are surrounded by microglia, phagocytic immune cells which participate in the clearance of Aβ (Hemonnot, Hua, Ulmann, & Hirbec, 2019; Mattiace, Davies, Yen, & Dickson, 1990). The ultrastructure of amyloid plaques has, so far, been studied by TEM (Gowrishankar et al., 2015; Terry, Gonatas, & Weiss, 1964) but also by correlated light microscopy and FIB-SEM (Blazquez-Llorca et al., 2017). While the latter approach provides exact targeting of single events, ATUM-FIB allows unbiased sampling of cellular interactions around plaques. For ATUM-FIB, we collected 18 consecutive sections at 5 µm thickness onto CNT tape. 3D reconstruction of the light and electron micrographs revealed the amyloid plaque distribution in the resulting 90 µm-thick volume, as well as the vasculature pattern and other morphological features (Fig. 5A,B).

**Figure 5.**
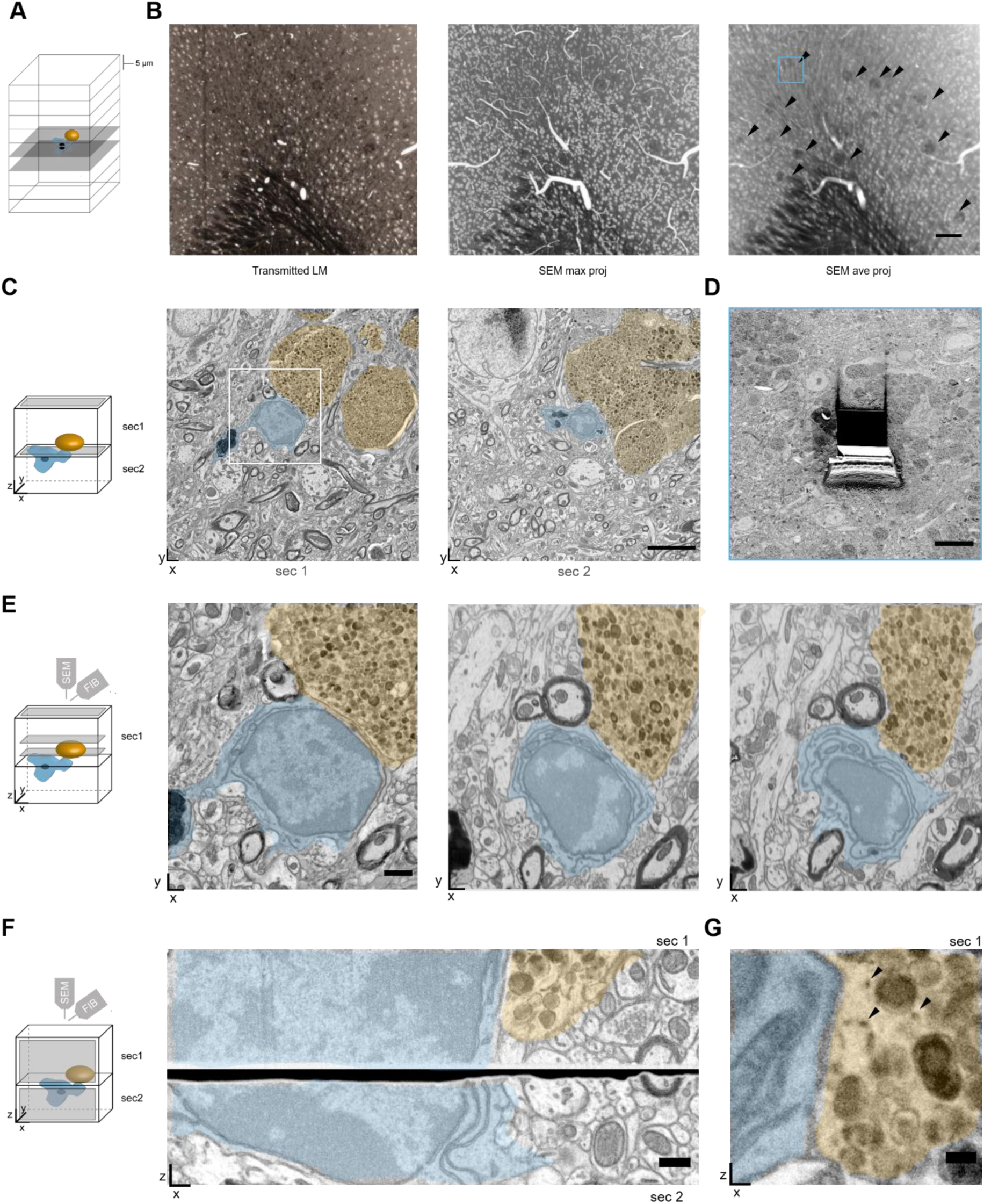
Targeted FIB-SEM on selected consecutive sites of semithick sections in FAD cortex samples. (A) Schematic of ATUM-FIB on consecutive 5 µm sections of mouse FAD cortex samples. Microglia (blue) contacts to plaques (orange) are targeted. (B) Transmitted light image of one section (20x objective, left). Maximum projection of 18 consecutive section BSD images reveals blood vessel and neuronal cell body distribution (middle). In the average projection thereof the plaque pattern (arrowheads) and the location of the target plaque (blue box) are highlighted. Scale bars 100 µm. (C, E, F, G) Schematics showing the imaging planes (grey) in the respective subfigures. (C) Two consecutive 5 µm semithick sections of the FAD cortex sample on CNT tape. The microglial cell (blue) and dystrophic neurites (orange) are highlighted. The white box indicates the magnified image in (E). Scale bar 5 µm. (D) Trench milled at the target site in section 1. (E) Magnified surface image of the microglia to plaque contact site of section 1 (left). This region was targeted for FIB-SEM at 5×5×5 nm. Reconstructions of two deeper levels within the semithick sections are shown (middle, right). Scale bar 1 µm. (F) Stitched cross sections of corresponding areas in consecutive semithick sections 1 and 2 after FIB-SEM. Scale bar 500 nm. (G) Regions of interest of selected sections of a FIB-SEM run (section 1) revealing ER structures (arrowheads) at sites of contact between microglia and dystrophic neurites. Scale bar 200 nm.

We selected two consecutive sections containing a plaque surrounded by a microglial cell for FIB-SEM examination (Fig. 5A-C). The region of interest for the deposition of a protective carbon layer was relocated by overlaying the SEM image from the section series with a BSD image acquired at 8 kV. We imaged a 20 × 20 × 5 µm volume at 5 × 5 × 5 nm by FIB-SEM (Fig. 5D). In order to reveal the strength of isotropic imaging we aligned the FIB-SEM volume and reconstructed different xz plane images of the microglial cell. The original SEM surface image was comparable to these virtual sections (Fig. 5E). Notably, we were able to stitch cross sections across consecutive sections (Fig. 5F), enabling extended z-volume analysis beyond 5 µm. Dystrophic neurites displayed ER structures at the contact sites opposing the microglial plasma membrane (Fig. 5G). While the target volume is resolved at high resolution and with isotropic voxels, light and electron microscopic images of the surrounding tissue provide a morphological context. Moreover, the non-destructive nature of serial sectioning on tape allows for reinspection of other regions of interest at high resolution.

## Discussion

Here, we introduce ATUM-FIB as a straightforward approach to combine overview imaging of tissue sections with targeted high-resolution three-dimensional reconstruction of subvolumes. In summary, ATUM-FIB has the following of advantages over previously developed 3D EM volume workflows (Fig. 6): (1) The ATUM-based generation of semithick sections allows screening of larger tissue volumes for rare events or specific sites of interest based on either light microscopic or low resolution SEM exploration of the semithin sections. Such sites can then be targeted by FIB-SEM with ∼5 nm isotropic xyz resolution. Thereby, the high resolution volumes are put into a larger morphological context, similar to microCT investigations (Handschuh, Baeumler, Schwaha, & Ruthensteiner, 2013; Starborg et al., 2019), but with the advantage of providing direct access to target structures instead of coordinates. Moreover, this tissue context provides rich fiducial landmarks for correlated light/ electron microscopy (Luckner et al., 2018). (2) In contrast to standard ultrathin ssTEM/SEM or ATUM techniques, our approach results in 100-fold fewer sections that need to be archived. This saves time, simplifies handling and reduces cost. (3) Just as ATUM, ATUM-SEM is a non-destructive technique (except for the FIB-targeted areas), and thus preserves a library of semithin sections. These can be revisited to increase the number of detailed observations or explore new questions that emerge over time. The limitation in volume depth of ATUM-FIB due to the maximum thickness of semithick sections (≤5 μm) compared to investigations of whole blocks, can be overcome be analysing the same region in consecutive semithick sections. Stitching of regions across serial sections has been successfully applied before for ultrathick partitioning (Kenneth J. Hayworth et al., 2015) as well as (S)TEM tomography (Aoyama, Takagi, Hirase, & Miyazawa, 2008; Baumeister, Grimm, & Walz, 1999; He & He, 2014), and has been shown to result in minimal loss of information (Kenneth J. Hayworth et al., 2015) (Suppl. Fig. 1). (4) In contrast to ultrathick partitioning (Kenneth J. Hayworth et al., 2015), surface SE/BSD information of consecutive semithick ATUM-FIB sections can be matched and aligned without the need FIB-SEM information of the containing tissue section. Moreover, there is no need to section into oil instead of standard water baths, which circumvents the need for re-embedding. (5) With the adaptation of standard diamond knives, our method can be applied in any EM lab with established FIB-SEM and ATUM workflows without the need of further equipment. This includes the standard rOTO staining method employed, circumventing the need to establish novel en bloc contrasting protocols.

**Figure 6.**
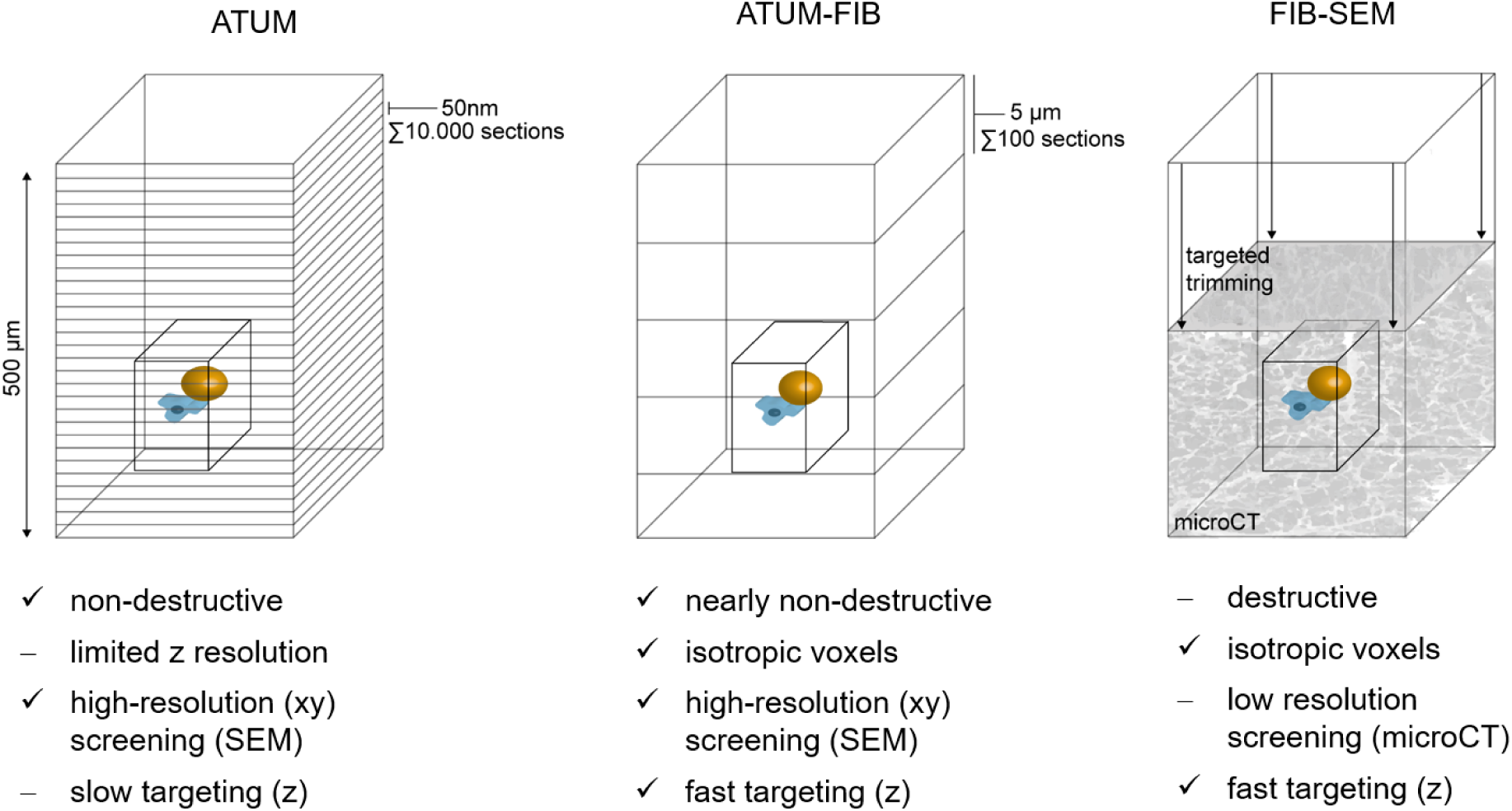
Comparison of volume SEM techniques for targeting rare events. Structures of interest like cell-cell interactions (microglia, magenta, dystrophic neurite, ocher) can be visualized by volume EM techniques. Current approaches for targeted imaging involve Automated Tape Collecting Ultramicrotomy (ATUM) and Focused Ion Beam Scanning EM (FIB-SEM). ATUM-FIB combines the advantages of both approaches. High-precision targeting based on ultrastructural features is enabled while increased section thickness reduces screening time and complexity compared to ATUM. Targeted FIB-SEM targeting is usually based on microCT data with limited xy resolution combined with trimming (arrows). A 500 µm z depth would be covered by 100 (at 5 mm thickness, ATUM-FIB) instead of 10.000 (at 50 nm thickness, ATUM) sections. Comparable to FIB-SEM, ATUM-FIB allows for acquisition of the region of interest (black box) at isotropic voxels.

Based on these advantages, we envisage a broad range of applications, as the implementation is comparably simple and imaging modalities and section thickness can be flexibly adapted to scientific questions and biological tissues of interest. The information content of subsequent isotropic FIB-SEM investigations is especially suited for the analysis of organellar ultrastructure and cell-cell contacts in defined physiological or pathological circumstances previously screened on serial semithick sections. As an example of such an application, we here demonstrate that ATUM-FIB can reveal the ultrastructure of cellular contact sites between microglia and axons in amyloid plaques in FAD mouse cortex. The fact that ATUM-FIB preserves the tissue library and provides histology-like context, makes the method especially attractive in settings, where rare and precious samples are being archived for long-term reinvestigations. This includes material from complex treatment studies or correlated in vivo imaging/EM investigations (Follain, Mercier, Osmani, Harlepp, & Goetz, 2017; Matthia A. Karreman et al., 2014) in animals, but especially also for multi-scale investigations of human samples, e.g. brain biopsies that require cross-referencing with standard histopathology - and will be increasingly needed to validate ultrastructural findings, e.g. from animal models of disease (Lewis et al., 2019) (Jonkman et al., 2019; Shahmoradian et al., 2019).

## Acknowledgements

This work was supported by DFG under Germany’s Excellence Strategy within the framework of the Munich Cluster for Systems Neurology (EXC 2145 SyNergy—ID 390857198) and the TRR 274/1 2020 (project Z01; ID 408885537) T.M.’s lab was also supported by DFG FOR2879, A03 and SFB/TRR274, projects B03, C02. We thank Katalin Völgyi and Ozgun Gokce for providing fixed brain samples, Richard Schalek, Mark Terasaki and Gerhard Wanner for valuable scientific and technical advice and Felix Beyer and Kerstin Karg for technical assistance.

## Author contribution

M.Sch., M.K., T.M., M.S. conceived the project, M.Sch. designed the experiments. G.K. and M.Sch. carried out experiments and analysed the data. H.G. adapted and provided the diamond knives. M.Sch. wrote the first draft and M.K., T.M., M.S. the final version of the manuscript.

---

**Supplementary Figure 1.**
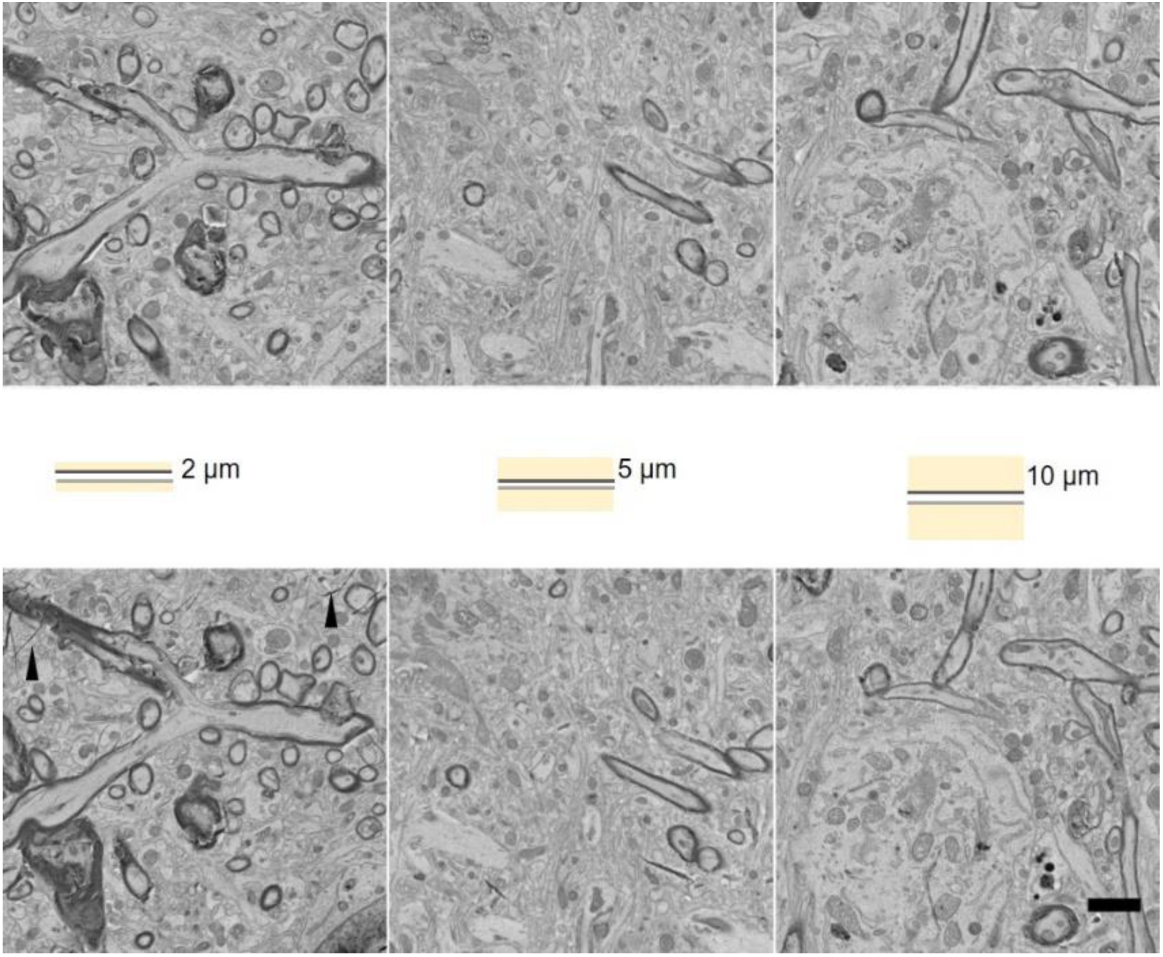
Consecutive semithick section tissue loss. (A) Schematics of opposing section surfaces revealing one section top face (light grey) and the bottom side of the consecutive one (dark grey). The remaining cross section thickness is shown in yellow. (B) Matching section BSD images of consecutive 2, 5 and 10 µm sections are shown. The first row shows the bottom of one section and the lower row the opposing surface of the following section. The latter were turned upside down and deposited on CNT tape. Folds (arrowhead) could not be prevented in this turning procedure of thinner sections (< 5µm). Scale bar 2 µm.

## Online Methods

### Sample preparation

FAD (2 months) and wt (18 months) mice were perfused with fixative containing 2.5% glutaraldehyde (Science Services), 2% PFA (Science Services) and 2 mM CaCl_2_ in 0.1 M sodium cacodylate buffer (Science Services). Brains were dissected and transferred into fixative for incubation for two additional days. Tissue sections of max. 1mm thickness comprising cortex and corpus callosum regions were prepared.

Fixed samples were stained *en bloc* by a varied Hua rOTO protocol (Hua et al., 2015; Tapia et al., 2012) without lead aspartate staining. We applied a sequence of reduced 2% osmium tetroxide in 0.1 M cacodylate buffer pH 7.4 followed by 2.5% potassium hexacyanoferrate in the same buffer. After washes the tissue was incubated in 1% aqueous thiocarbohydrazide (TCH) and subsequently in 2% aqueous osmium tetroxide. After overnight incubation in 1% uranylacetate at 4°C and 2h in 50°C, samples were dehydrated and infiltrated at least 2h at different resin in acetone concentrations (25, 50, 75%) and overnight and for another 4h in 100% of the respective resin. Durcupan (Science Services) resin was prepared by mixing 11.4 g of component A (epoxy resin), 10.0 g of component B (964 hardener), 0.1 mL of component D (dibutyl phthalate) for the infiltration steps and the same mixture including component C (964 accelerator). For standard epon (Serva) 21.4 g glycidether 100 with 14.4 g dodecenylsuccinic anhydride (DDSA) and 11.3 g nadic methyl anhydride (NMA) were combined for 10 min and 0.84 mL 2,4,6 tris(dimethylaminomethyl)phenol (DMP-30) were added while stirring for another 20 min. LX112 resin (LADD) was prepared by mixing 4.5 g of mix A (mixture of 5 g of LX112 and 6,45 g of nonenyl succinic anhydride), 10.5 g of mix B (mixture of 5 g of LX112 and 4.35 g of NMA) and 0.6 ml of DMP-30 (all components from LADD research industries). Resins were cured at 60 °C for 10h, 15h or 48h.

### Automated tape-collection ultramicrotomy (ATUM)

Epon blocks were roughly trimmed with EM TRIM2 (Leica) and subsequently, a rectangular tissue block (∼ 2×1.5×0.2 µm) was exposed using the trimtool 45 diamond knife (Diatome). Thick sections were initially generated using a histo jumbo knife (45°, 6 mm, Diatome) in a RMC ultramicrotome (Powertome). The 35° and 45° ultra knife boats for the custom-made heated knives were provided by Diatome. Two holes (3 and 6 mm diameter, respectively) were milled into the base part of the knife boat to fit a temperature sensor (cable probe 3×30 mm, Sensorshop) and a heater (Hotend Heater Catridge CNC for 3D printer, 24V, 40W; Sensorshop). We used a digital on/off temperature regulator (for PT100, Sensorhop). Standard infrared lights (230V, 150W, Conrad Electronics) were installed at both sides of the microtome. Temperature was controlled by standard probe (VWR) and infrared thermometers (Conrad Electronics) to values within the range of 35-45°C.

Single sections were fished from the water bath by ∼0.3×0.8 mm carbon nanotube tape (CNT) tape (Science Services) pieces using inverse forceps. For serial sectioning the RMC tape collector was adapted to the histo jumbo knife by bypassing the tension lever and guiding the CNT tape behind the knife directly to the collector nose. The spatial arrangement of the tape collector nose had to be adapted to the heating unit. Sectioning speed was set to 0.2-0.4 mm/sec with increased tape speed within (0.4 mm/sec) and reduced tape speed outside (0.1 mm/sec) the cutting window. This assured efficient uptake and minimization of empty intersection space on the tape. Slow speed was needed both for limiting compression, as well as for keeping the sections longer in the heated water bath to smoothen. If needed, sections were guided onto the tape collector using fine brushes.

### Slide scanner serial light microscopy

For light microscopic investigation, CNT tape strips with single sections or 5 cm strips with serial sections were positioned on a glass slide. For better adherence, a few drops of water were placed between tape and glass placed on a heating plate (80°C). Good adherence was important for tape flattening as a prerequisite of the slide scanner autofocusing function. Serial light microscopy was performed on a slide scanner (Pannoramic MIDI II 2.0.5, 3D Histech). We selected sections by thresholding and imaged using the autofocusing and the extended focus level (9 focus levels, focus step size 0.2 µm x 5) functions using the 20x objectives. By choosing the extended focus option, the software selects the sharpest image from each focus level for each image field, and combines them into one single image. The autofocus was restricted and the range set by testing it for several sections on different slide locations. Jpeg files were generated from the original data using the Pannoramic software CaseViewer2.2 (3D Histech).

### Serial scanning electron microscopy

Sections on glass slides were postcured for 30-48h at 60°C. CNT strips with tissue sections were detached and assembled onto carbon tape (Science Services), mounted onto a 4-inch silicon wafer (Siegert Wafer) and grounded with adhesive carbon tape. Serial section images were acquired on a Crossbeam Gemini 340 SEM (Zeiss) in backscatter mode at 4 keV (high gain) at 7-8 mm WD and 30 or 60 µm aperture. In ATLAS5 Array Tomography (Fibics, Ottawa, Canada) a wafer overview map at 1000-3000 nm/pixel was generated. On this basis, sections were mapped and imaged at medium (60 × 60 – 100 × 100 nm) resolution. Regions of interest from these section sets were acquired at 10×10 nm/pixel (2 µs dwell time, line average 2). Image series were aligned in TrakEM2 using a combination of automated and manual processing, registered and analysed in Fiji (Schindelin et al., 2012).

### Focused Ion Beam Scanning Electron Microscopy (FIB-SEM)

Selected thick sections on CNT tape were cut from the silicon wafer including the adhesive carbon tape underneath using a scalpel. Theses samples were mounted with conductive carbon cement (LEIT-C, Plano) and conductive silver colloid (Plano) onto standard aluminum stubs (Plano). A thin layer of carbon was sputtered onto the sections (carbon cord, Q150T ES, Quorum). Milling and imaging were performed on a Crossbeam Gemini 340 FIB-SEM operating under SmartSEM (Zeiss) and Atlas-3D (Fibics Incorporated). Ion beam currents of 50 pA - 15 nA were used. The milling rate was set to 5 nm slices. SEM images were recorded with an aperture of 60 μm in the high current mode at 2 kV of the InlenseDuo detector with the BSE grid set to 300-500 V and the SE detector. Voxel sizes of 5 × 5 × 5 nm were chosen. Images series of 1000-2000 consecutive sections were recorded. In ATLAS, the milling current and depth were adjusted to match with exposure time of the SEM (line average 2, dwell time 3 µs). Automatic correction of focus (auto tune) and astigmatism (auto stig) was applied every 30 minutes. FIB-SEM image stacks were aligned and analyzed in Fiji (Schindelin et al., 2012).

